# dsRNA-Seq: Identification of viral infection by purifying and sequencing dsRNA

**DOI:** 10.1101/738377

**Authors:** Carolyn J. Decker, Halley R. Steiner, Laura L. Hoon-Hanks, James H. Morrison, Kelsey C. Haist, Alex C. Stabell, Eric M. Poeschla, Thomas E. Morrison, Mark D. Stenglein, Sara L. Sawyer, Roy Parker

**Affiliations:** Department of Biochemistry, University of Colorado, Boulder, CO, USA; Howard Hughes Medical Institute, Chevy Chase, MD, USA; Department of Microbiology, Immunology, and Pathology, Colorado State University, Fort Collins, CO, USA; Division of Infectious Diseases, University of Colorado School of Medicine, Aurora, CO, USA; Department of Immunology and Microbiology, University of Colorado School of Medicine, Aurora, CO, USA; Department of Molecular, Cellular and Developmental Biology, University of Colorado, Boulder, CO, USA

**Keywords:** dsRNA, double-stranded RNA, emerging disease, emerging viruses, RNA virus, RNA-Seq

## Abstract

RNA viruses are a major source of emerging and re-emerging infectious diseases around the world. We developed a method to identify RNA viruses that is based on the fact that all RNA viruses produce dsRNA while replicating. Purifying and sequencing dsRNA from total RNA isolated from infected tissue allowed us to recover replicating viral sequences. We refer to this approach as dsRNA-Seq. By assembling dsRNA sequences into contigs we identified full length RNA viruses of varying genome types infecting mammalian culture samples, identified a known viral disease agent in laboratory infected mice, and successfully detected naturally occurring RNA viral infections in reptiles. Here we show that dsRNA-Seq is a preferable method for identifying viruses in organisms that don’t have sequenced genomes and/or commercially available rRNA depletion reagents. Similar to other metagenomic strategies, dsRNA-Seq has the potential to identify unknown viral disease agents that share little to no similarity to known viruses. However, the significant advantage of this method is the ability to identify replicated viral sequences, which is useful for distinguishing infectious viral agents from potential noninfectious viral particles or contaminants.

## 1. Introduction

RNA viruses have a significant impact on human health and constitute a major source of emerging or re-emerging infectious diseases [1] such as MERS, Ebola, West Nile, Zika and chikungunya [2-3]. RNA viruses also pose a threat to animal and plant health, where they can cause major loss to crop and animal production and impact biodiversity [4-6]. In order to curb emerging infectious agents, it is useful to have robust methods to identify new viruses. Additionally, the threat of synthetic viruses [7] presents serious challenges to modern approaches to viral identification since those viruses need not share similarity to previously known viral agents. It is therefore useful to have multiple effective means to identify new viral disease agents.

High-throughput sequencing of total RNA isolated from infected individuals is a powerful approach to identifying and determining complete genomes of RNA viruses that are potentially responsible for diseases without a known causative agent. In addition, this approach allows for the detection of variants of known viruses or synthetic viral agents, which may not be recognized by PCR or serological based techniques. One limitation of this strategy is that viral sequences are present at very low levels relative to host sequences in clinical samples, which limits the sensitivity of viral detection and the ability to reconstruct viral genomes [8-11].

There are several current approaches that enrich for viral sequences to improve the sensitivity of detection. Because the vast majority of host RNA is ribosomal RNA, one approach has been to remove host rRNA sequences [9,12]. Another approach is to enrich for viral particles from infected samples [13], the feasibility of which is dependent on the sample type and whether sufficient viral particles can be obtained. Positive selection strategies have also been developed wherein viral sequences are “captured” by hybridization to virus-specific probe sets based on known viruses [14-16]. However, these strategies may be biased against detecting novel viruses depending on how closely they are related to known viruses.

We have developed an alternative method to enrich for viral RNA sequences that incorporates both negative selection to remove host RNA and positive selection for replicating viruses that is not based on known viral sequences. We reasoned that all RNA viruses produce double stranded RNA (dsRNA) while replicating, therefore by purifying dsRNA from total RNA isolated from infected tissue we would enrich for RNA viral sequences. This approach would also allow us to distinguish nonreplicating viral particles from replicating infectious agents. To purify dsRNA, we first remove the majority of host RNA by treating with a single-strand specific RNase, and then isolate the dsRNA by immunoprecipitation with a sequence-independent anti-dsRNA antibody [17-18]. We then sequence and de novo assemble the resulting dsRNA to aid in viral discovery. We refer to this approach as dsRNA-Seq and have successfully used it to identify a variety of replicating RNA viruses in infected animal tissues.

## 2. Materials and Methods

### 2.1. dsRNA purification

Total RNA from Vero cells infected with dengue virus type 2 (New Guinea C), influenza A (H3N2 Udorn) or mock-infected was extracted using Trizol (Thermo-Scientific) [19] and provided by Alex Stabell and Sara Sawyer (University of Colorado, Boulder). Total RNA extracted from whole quad muscle collected 5 days post-infection from C57BL/6J mice that were mock-infected or infected with Ross River virus (T48) as described in [20] was provided by Kelsey Haist and Thomas Morrison (University of Colorado, Anschutz Medical Campus). All mouse studies were performed at the University of Colorado Anschutz Medical Campus (Animal Welfare Assurance #A 3269-01) using protocols approved by the University of Colorado Institutional Animal Care and Use Committee and in accordance with the recommendations in the Guide for the Care and Use of Laboratory Animals of the National Institutes of Health. Total RNA extracted from green tree python lung and pooled lung/esophagus, rough-scaled python lung, mule deer brain and lymph node, and boa constrictor kidney tissue as described in [21] was provided by Laura Hoon-Hanks and Mark Stenglein (Colorado State University). All of the reptile and deer samples were collected postmortem from client-owned animals for diagnostic assessment. dsRNA was purified from 100μg total RNA isolated from Vero cells, 5μg total RNA from mouse skeletal muscle samples, and 10μg total RNA from reptilian and mule deer samples. Total RNA, at final concentration of 0.2μg/μl, was incubated with 1 unit RNase 1 (Ambion) (10 units RNase 1 for reptilian and mule deer samples) and 0.2 units Turbo DNase1 (Ambion) per μg total RNA in 1X Turbo DNase 1 buffer (which contains 75mM monovalent salt) and 125mM NaCl (final monovalent salt concentration 0.2M) at 37°C for 30 min. Reactions were then diluted with buffer pre-chilled on ice to final concentration of 20mM TrisCl pH 7.5, 0.15M NaCl, 0.2mM EDTA, 0.2% Tween20 to final volume of 500μl for skeletal muscle samples or 1ml for Vero cell samples and then incubated with 5μg of J2 anti-dsRNA antibody (Scicons) [17] pre-bound to 0.75μg of Protein A Dynabeads (Invitrogen) with end to end rotation at 4°C for 2hr. Beads were recovered using a magnet and washed 3 times with 1X IP buffer (20mM TrisCl pH 7.5, 0.15M NaCl, 0.1mM EDTA, 0.1% Tween20). The dsRNA was recovered from the beads by adding 150μl 1X IP buffer and 450μl of Trizol LS (Ambion) and then following the manufacturer’s protocol for isolating RNA. The resulting aqueous phase was mixed with equal volume 70% ethanol, applied to RNA Clean-Up and Concentration Micro-Elute columns (Norgen Biotek) following manufacturer’s protocol and the dsRNA eluted in 15μl of nuclease free water.

### 2.2. RNA library construction and sequencing

For Vero cell culture samples, 9μl dsRNA in water was denatured at 95°C 2min then cooled on ice, RNA libraries were then prepared using ScriptSeq v2 Stranded Kit (Epicentre) following the manufacturer’s protocol except fragmentation was done at 85°C for 8min. Each sample was indexed with 6 bp unique barcode, libraries were pooled and sequenced on MiSeq (75base PE reads, Illumina). For mouse, reptilian and mule deer tissue samples, 11μl dsRNA diluted to final 10mM TrisCl pH 8.0, 0.1% Tween 20 in 12μl was denatured at 95°C 2min then cooled on ice, RNA libraries were then prepared using Ovation SoLo RNA-Seq System (NuGen) starting at Step C and stopping after Step L (Library Amplification I Purification) in manufacturer’s protocol therefore the dsRNA libraries did not undergo the ribosomal sequence depletion steps in the protocol. RNA libraries were also prepared from 2ng of total RNA from the mouse samples using Ovation SoLo RNA-Seq System (NuGen) following the manufacturer’s protocol starting at Step C, including the ribosomal sequence depletion steps and stopping after Step P (Library Amplification II purification). Each sample was indexed with 8 bp unique barcode, libraries were pooled and sequenced on NextSeq (75base PE reads, Illumina) using SoLo Custom R1 primer and standard Illumina R2 primer.

### 2.3. Analysis of dsRNA-Seq sequences from Vero cell samples

Illumina adaptors were trimmed in PE mode, reads trimmed when average quality score in 4 base window fell below 20 and then reads smaller than 50nt were discarded using Trimmomatic 0.32 [22]. Reads were assembled into contigs 500nt or longer using Trinity 2.0.6 [23] and the longest isoform of related contigs was selected using the Trinity helper script get_longest_isoform_seq_per_trinity_gene.pl. Contigs were mapped to the *Chlorocebus sabaeus* genome (GCF_000409795.2), dengue virus type 2 (NC_001474.2) and influenza A virus (NC_007366.1) genomes using bwa-mem algorithm of bwa 0.7.15 [24]. To look for similarity between the contigs and known sequences the contigs were used as queries using BLASTn command of BLAST 2.7.1 [25] against the NCBI nt database. Contigs with similarity to phiX174 were removed. Bowtie 2.2.9 [26] was used to map the dsRNA-Seq reads to the assembled contigs, *Chlorocebus sabaeus*, dengue virus type 2 and influenza A virus genomes. To determine which contigs were derived from dsRNA, reads that were derived from the forward strand of each contig were extracted using SAMtools 1.8 [27] samtools view and flag options -f 64 -F16 for Read 1 and -f128 -F16 for Read 2. The number of forward strand reads and the total number of reads that mapped to each contig was then determined using samtools idxstats and used to calculate the percentage of forward strand reads. The strand-specificity of Scriptseq libraries is >98% (Epicentre product literature) therefore contigs with < 98% of forward strand reads were considered being derived from dsRNA rather than possible contaminating ssRNA.

### 2.4. Analysis of dsRNA-Seq and ribodepleted RNA-Seq sequences from mouse skeletal muscle samples

Adaptor and additional sequences flanking library inserts were removed using BBduk.sh of bbmap 38.05 [28] available at https://sourceforge.net/projects/bbmap/), reads were trimmed when average quality score in 4 base window fell below 20 and reads smaller than 50nt were discarded using Trimmomatic 0.36 [22]. Processed dsRNA-Seq reads were mapped to GRCm38 mouse genome using Bowtie 2.2.9 [26]. The fastq command with -f 12 option of SAMtools 1.8 [27] was used to extract matched paired reads where both members of the pair did not map to the mouse genome. The reads that did not map to the mouse genome were assembled into contigs at least 750 nt long using Trinity 2.6.6 [23], the longest isoform of related contigs was selected and which contigs were derived from dsRNA was determined as described for the Vero samples except the strand-specificity of Ovation SoLo libraries is >90% (NuGen product literature) therefore contigs with < 90% of forward strand reads where considered being derived from dsRNA rather than possible contaminating ssRNA. To look for similarity between the dsRNA contigs and known sequences the contigs were used as queries using BLASTn command of BLAST 2.7.1 [25] against the NCBI nt database. Contigs from each dataset were mapped to Ross River virus strain T48 (GQ433359.1) using bwa mem of bwa 0.7.15 [24]. The dsRNA-Seq and ribo-depleted RNA seq reads that aligned to the Ross River virus strain T48 genome were identified using Bowtie 2.2.9 and the –no-unal option. Note, the 63nt poly A tail at the 3’ end of the Ross River virus was deleted for this analysis. Reads that were derived from the positive strand of Ross River virus were extracted using SAMtools 1.3.1 samtools view and flag options -f 64 -F16 for Read 1 and -f128 -F16 for Read 2. Reads derived from the negative strand with flag -f 80 for Read 1 and -f 144 for Read 2. The number of positive strand reads, negative strand reads and total number of reads was then determined using samtools idxstats and bedgraphs were produced using genomeCoverageBed command of BEDTools 2.25.0 [29]. A similar strategy was used to analyze dsRNA-Seq and ribo-depleted RNA-Seq read mapping to the mouse mitochondrial chromosome.

### 2.5. Analysis of dsRNA-Seq sequences from reptile and deer tissue samples

We processed the snake and deer dsRNA-Seq samples, assembled reads into contigs and determined which contigs were derived from dsRNA as described for the mouse samples. To investigate similarity between the dsRNA contigs and known sequences, the contigs were used as queries using BLASTn against the entire NCBI nt database, BLASTx againt nr, and BLASTx limited to viral sequences. We used the same mapping strategy as in the mouse sample analysis in order to assemble partial and full viral genomes; we mapped the dsRNA contigs back to viral genomes previously found using standard RNA-seq and to genomes of any new viruses discovered and displayed the alignment in IGV. If contigs only hit to viral proteins, we used sequences as queries in BLASTx and recorded the subject coverage and identity percentage to determine where these contigs were aligning to on the viral genome. RNA-Seq libraries and data analysis for the boa constrictor and chameleon samples were generated as previously described [21].

### 2.6. Sequence Data Availability

dsRNA and total RNA sequence data are available as raw reads from the NCBI Short Read Archive (SRA) under study accession number SRP201404. Individual accession numbers are Vero_1_dsRNA, SRR9301166; Vero_2_dsRNA, SRR9301167; Vero_3_dsRNA, SRR9301168; Mouse_1_dsRNA, SRR9301169; Mouse_2_dsRNA, SRR9301170; Mouse_3_dsRNA, SRR9301171; Mouse_4_dsRNA, SRR9301172; Mouse_5_dsRNA, SRR9301173; Mouse_1_RNA, SRR9301164; Mouse_2_RNA, SRR9301165; Mouse_3_RNA, SRR9301160; Mouse_4_RNA, SRR9301161; Mouse_5_RNA, SRR9301162; Green_Tree_Python _lung_dsRNA, SRR9301163;

Green_Tree_Python_lung_esophagus_dsRNA, SRR9301156; Rough_Scaled_Python_lung_dsRNA, SRR9301157; Boa_Constrictor_kidney_dsRNA, SRR9301158; Veiled_Chameleon_lung_trachea_oral mucosa_dsRNA, SRR9301159; Veiled_Chameleon_lung_liver_kidney_1_dsRNA, SRR9301154; Veiled_Chameleon_lung_liver_kidney_2_dsRNA, SRR9301155;

Mule_Deer_brain_dsRNA, SRR9301151; Mule_Deer_lymph_node_dsRNA, SRR9301150; negative_control, SRR9301153; Boa_Constrictor_kidney_RNA, SRR9301152.

## 3. Results

### 3.1. dsRNA-Seq detects viral infections of cultured mammalian cells

As an initial test to determine whether we could detect viral infections in mammalian cells by purifying and sequencing dsRNA, Vero cells were infected with influenza A virus, dengue virus type 2, or were mock infected. Total RNA was isolated from the mock and infected cells and samples were blinded for the remainder of library preparation, sequencing, and data analysis.

dsRNA was purified from the total RNA using a two-step protocol (Figure 1A, see Materials and Methods). First, the total RNA was treated with DNase 1 and a single-strand specific RNase to remove any contaminating DNA and to enrich for double-stranded RNA. The RNase treatment was performed in the presence of 0.2 M monovalent salt to stabilize base pairing interactions to minimize the inadvertent digestion of dsRNA. Subsequently, an antibody that recognizes dsRNA [17-18] was used to immuno-purify the dsRNA. The anti-dsRNA antibody is highly specific for dsRNA, requires at least 40 base pairs of dsRNA for binding, and is sequence-independent, although it does have some preference for binding particular AU rich sequences [17-18]. Testing this approach using total RNA spiked with varying amounts of in vitro transcribed dsRNA revealed that 1) dsRNA was specifically enriched over single-stranded RNA and 2) the enrichment of the dsRNA was dependent on the anti-dsRNA antibody (Figure S1). Moreover, a broad range of dsRNA can be efficiently isolated. We observed 50-100% recovery of 100ng to 10pg of dsRNA (Figure S1).

**Figure 1.**
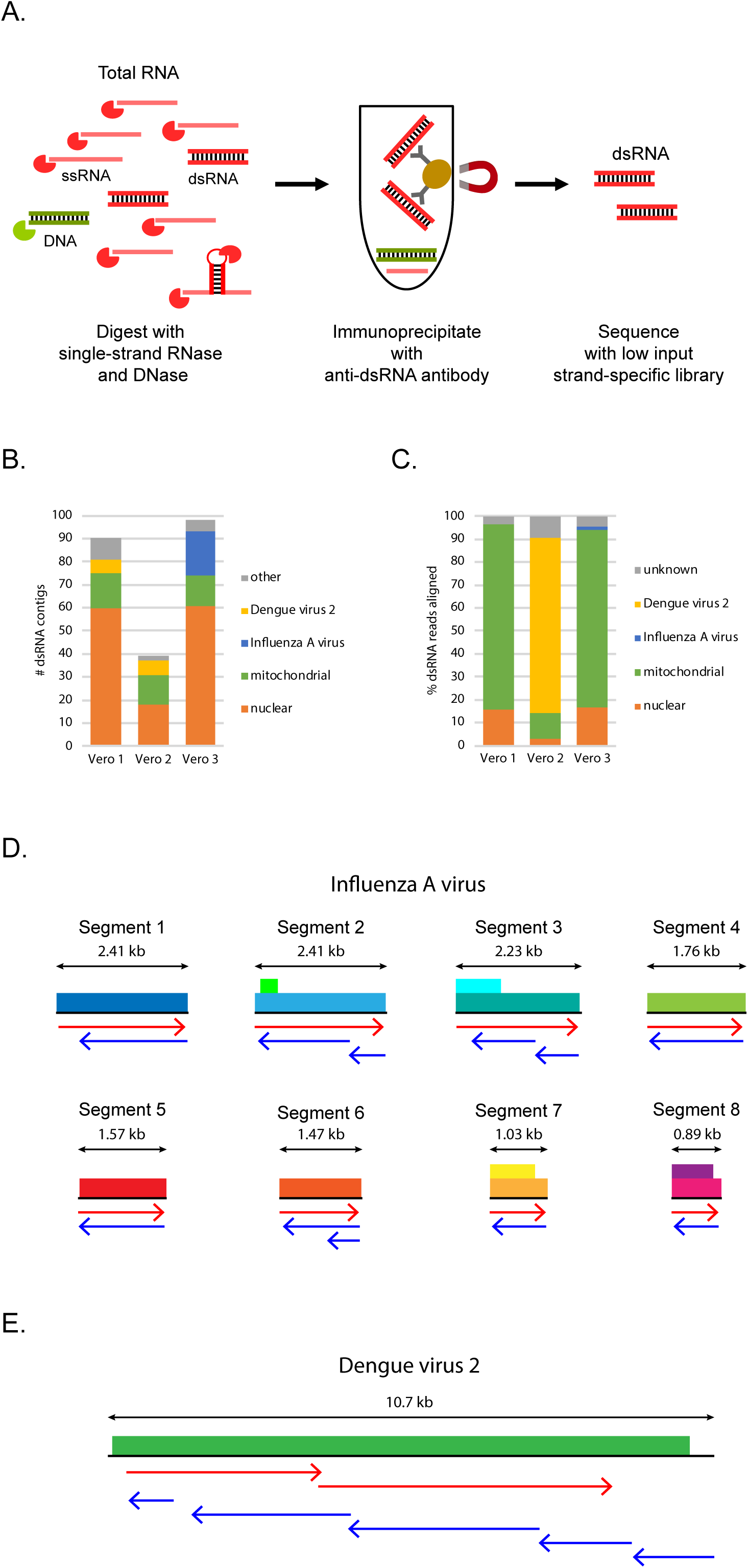
dsRNA-Seq detects viral infections of cultured mammalian cells. (a) Outline of dsRNA purification method; (b) Number of dsRNA contigs assembled from dsRNA-Seq reads from infected or mock infected Vero cell samples and their classification based on mapping to host nuclear or mitochondrial chromosomes or BLASTn analysis against NCBI nt; (c). Percentage of dsRNA-Seq reads that align to the host nuclear or mitochondrial chromosomes, influenza A viral genome, dengue virus type 2 genome or did not align (unknown). For (d) and (e), viral genomes are illustrated with protein coding regions indicated by colored boxes. Arrows indicate alignment of contigs to viral genomes or genome segments. Contigs representing the positive strand are in red; negative strand in blue; (d) Alignment of contigs assembled from Vero 3 sample to influenza A viral segments; (e) Alignment of contigs assembled from Vero 2 sample to dengue virus type 2 genome.

dsRNA was isolated from the Vero cell total RNA using the two-step purifation scheme and low input cDNA libraries were prepared from the dsRNA and sequenced. All libraries sequenced were constructed to maintain strand-specific information, which allows us to determine if a sequence detected was present as dsRNA (see Materials and Methods and below).

To mimic a situation where we were trying to identify a virus of unknown sequence, we first assembled the short dsRNA reads into longer contigs (see Materials and Methods) in an attempt to assemble virus genomes. From each of the Vero cell samples we assembled between 39-98 contigs that were 500 bases or longer (Figure 1B and Table S1). Because all three samples, including the mock-infected sample, had similar numbers of contigs, we reasoned that many contigs may be derived from host dsRNA rather than viral dsRNA. Mapping the contigs to the *Chlorocebus sabaeus* (green monkey) genome revealed that the majority of the contigs aligned to the host nuclear or mitochondrial chromosomes (Figure 1B). One possibility is that the host contigs were derived from contaminating single-stranded transcripts. However, when we mapped the reads to the contigs we found that both strands of each contig were represented in the dsRNA reads (see Materials and Methods), consistent with the contigs being derived from host dsRNA (Table S1). Mapping the dsRNA reads to the *Chlorocebus sabaeus* genome revealed that in two of the samples over 90% of the reads are in fact from the host, with the vast majority of the dsRNA coming from the mitochondrial genome (Figure 1C). Thus, dsRNA-Seq reveals the presence of sense/antisense and/or dsRNA in mammalian cells.

To determine if any of the remaining contigs were similar to known viruses, we used BLASTn to look for similarities between the nucleotide sequence of the contigs and sequences in the NCBI nucleotide database. In the Vero 3 sample, nineteen of the contigs shared significant similarity to influenza A virus (Figure 1B and Table S1). In contrast, the other two samples did not have any contigs with similarity to influenza A virus (Figure 1B and Table S1). Influenza A is a negative sense single-stranded RNA virus composed of 8 segments. The nineteen contigs in the Vero 3 sample ranged in length between 0.5-2.3kb and represented both strands of all 8 segments of the virus (Figure 1D). Despite only 1.47% of the reads in the Vero 3 sample aligning to the influenza A virus (Figure 1C and Table S1), the entire genome was assembled, indicating that it is possible to detect viral infection using dsRNA-Seq even under conditions where the viral RNA is not highly represented in the recovered dsRNA.

In the Vero 2 sample, six contigs had BLASTn hits to dengue virus type 2 (Figure 1B and Table S1), a positive sense single-stranded RNA virus with a ∼11 kb monopartite genome. These contigs ranged in size from 0.8-9.9 kb and represented the entire dengue virus type 2 genome in both orientations except the 5’ most 300 nucleotides (Figure 1E). Dengue virus type 2 sequences were very abundant in the dsRNA isolated from the Vero 2 sample with 76% of the reads mapping to the dengue virus type 2 genome (Figure 1C and Table S1).

In contrast to the other two samples, no full-length or nearly full-length viral contigs were detected in the Vero 1 sample, which is consistent with this sample coming from the mock-infected cells. Although six contigs in the Vero 1 sample did share similarity to dengue virus type 2, these contigs were only 0.5-0.6 kb and did not cover the entire genome. Moreover, only 0.02% of the Vero 1 dsRNA mapped to the dengue virus type 2 genome (Figure 1C and Table S1) suggesting that these reads may have resulted from contamination from the Vero 2 sample, or by reads miss-assigned due to “index hopping” during sequencing [30].

Uncoding the samples revealed that the infectious agents were accurately identified by dsRNA-Seq, indicating that enrichment and sequencing of dsRNA is sufficient for the detection of positive and negative sense single-stranded RNA viruses in infected tissue culture cells.

### 3.2. dsRNA-Seq correctly detects viral infection in infected mice

We then asked if we could detect viral infections in animals by isolating and sequencing dsRNA from infected mouse tissue. Viral infection in infected animals or humans could be more challenging given that not every cell in a tissue will be infected, the viral load in infected cells may be low, and the amount of tissue in clinical samples may limit the amount of dsRNA that can be recovered and sequenced.

We obtained five samples of total RNA isolated from infected or uninfected mice. We were blinded to the number of the samples that were infected vs uninfected, the type of virus(es) used for infection, and the tissue from which the total RNA was isolated. Using western blot analysis with the anti-dsRNA antibody, we estimated the amount of dsRNA in the samples to be only ∼10-120 pg per 1 μg of total RNA (Figure S2). dsRNA was isolated from 5 μg of total RNA (∼50-600 pg dsRNA) from each sample and sequenced.

Since the tissue culture experiment suggested that the majority of dsRNA reads would be from the host, we first mapped the dsRNA sequences to the mouse genome. In each dataset, between 78 to 88% of the reads aligned to the mouse genome (Figure 2A, Table S2). In a second step, the reads that did not map to mouse sequences were assembled into contigs of 750 bases or longer in order to attempt to assemble full-length viral genomes or genome segments. We then used the strand-specific information in our library preparation to determine whether both strands of each putative dsRNA contig were represented in the dsRNA reads. This allowed us to distinguish between contigs derived from dsRNA versus from possible contaminating single-strand RNA. Four to sixteen contigs derived from dsRNA were assembled in each dataset and they ranged in length from 0.76 to 23 kb (Table S2).

**Figure 2.**
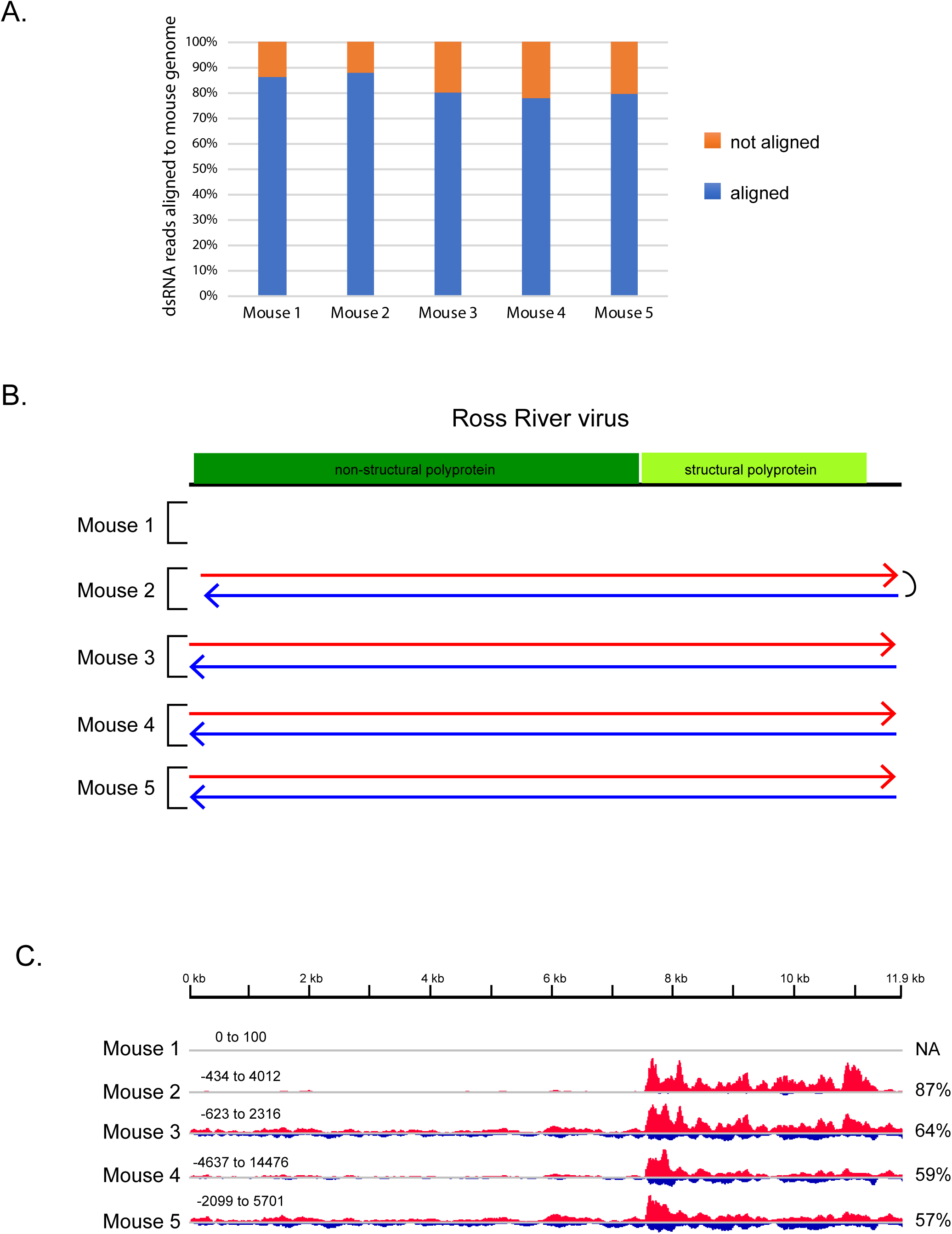
dsRNA-Seq detects viral infection in infected mice. (a) Percent dsRNA-Seq reads that aligned to the mouse genome; (b) Ross River virus with protein coding regions indicated in colored boxes. Arrows indicate the alignment of contigs assembled from dsRNA-Seq reads from mouse samples. Contigs representing the positive strand of the virus are in red; negative strand in blue. Mouse 2 sample contained a single contig that represented both the positive and negative strand of the virus indicated by the link on the right; (c) Bedgraphs of dsRNA-Seq reads that aligned to Ross River virus genome from mouse samples. Height indicates base count at each position along the Ross River virus sequence. Red indicates counts on positive strand. Blue indicates counts on negative strand. The percentage of the total reads that mapped to the Ross River virus that aligned to the positive strand is indicated on the right. Note: The bedgraphs have not been scaled relative to total number of dsRNA-Seq reads in each sample. Each read set was scaled individually, scale is indicated at top left for each set.

To understand the nature of these dsRNA contigs, we asked if they shared similarity to known viral or cellular sequences in the NCBI nucleotide database using BLASTn. We found that four of the five samples had contigs that were greater than 99% identical on the nucleotide level to Ross River virus (Figure 2B and Table S2). Ross River virus is an alphavirus-a single-stranded positive-sense RNA virus with a single genome segment of ∼11.9 kb. The contigs with similarity to Ross River virus represented full-length or nearly full-length viral sequences (Figure 2B). Uncoding the samples revealed that the four RNA samples with dsRNA contigs derived from Ross River virus came from skeletal muscle tissue of mice at 5 days post-infection with Ross River virus (T48 strain), with the remaining sample coming from a control mock-infected animal. Thus, dsRNA-Seq correctly identified the virus used for infection and differentiated between infected and non-infected animals.

Given that single-stranded RNA viruses which have not replicated would not be present as dsRNA, dsRNA-Seq should specifically detect ssRNA viruses that are or have replicated. Consistent with this idea, both positive-sense and negative-sense strands of Ross River virus RNA were well represented (Figure 2C) in the dsRNA-Seq datasets with 13-43% of the reads that aligned to Ross River virus being derived from the negative strand. Moreover, the high coverage of the negative strand allowed for the assembly of full length or nearly full length contigs representing the negative strand of Ross River virus from the infected animals (Figure 2B). Thus, the dsRNA-Seq analysis provided evidence of virus replication in the mouse tissue.

Synthesis of the negative strand of alphaviruses occurs early in infection followed by a switch to using the negative strand as a template to synthesize full-length genomic RNA as well as a subgenomic mRNA that encodes the viral structural proteins [31-32]. The mouse samples were collected 5 days post infection, past the peak of viral replication, therefore dsRNA-Seq is likely detecting negative strands produced earlier in infection. The abundance of reads mapping to the 3’ region of the virus (Figure 2C) is likely due to the high expression of the subgenomic RNA (see below and Discussion) which is synthesized at much higher levels than the full-length genomic RNA [33-34].

Additional contigs were assembled from the dsRNA-Seq reads. The majority of the remaining dsRNA contigs were derived from mouse sequences, primarily from mitochondria, or rRNA sequences from various organisms (Table S2). Contigs that represent single-stranded sequences (contigs for which reads only mapped to one strand) were also assembled from the reads (Table S2). Many of these contigs represent sequences from bacteria that are common contaminants in next generation sequencing libraries (Table S2) [35], which highlights the need to implement strategies to identify such contaminants, particularly when only small amounts of input RNA are available for library preparation [36].

We also identified single-stranded contigs that appear to be derived from viruses. For example, in all the mouse samples, we assembled contigs that were nearly identical to each other on the nucleotide level (Figure S3A) and encode proteins with similarity to picorna-like RNA viruses (Figure S3B and Table S2). These contigs are most likely contaminants of the dsRNA-Seq libraries rather than viruses that are infecting the mouse samples given that only their positive strand was represented in the dsRNA-Seq reads (Figure S3C). Moreover, reads corresponding to these viruses were not found in ribo-depleted RNA libraries prepared from the same total RNA samples (see below). Thus, an advantage of dsRNA-Seq libraries is the ability to distinguish between viral contigs that likely represent contaminating viral sequences and viral sequences of interest that are replicating.

### 3.3. Impact of dsRNA-Seq on detecting RNA viral infections

An unanswered question is how dsRNA-Seq compares to sequencing total ribo-depleted RNA for the identification of viruses. To compare the two approaches, we prepared ribo-depleted RNA-Seq libraries from the mouse tissue samples. We first compared the sensitivity of the two methods in detecting the Ross River virus. The percentage of reads that mapped to the Ross River viral genome was decreased in the dsRNA read datasets compared to the ribo-depleted RNA reads (Figure 3A). The percentage of Ross River virus reads was between 4-12 fold higher in the traditional RNA-Seq reads than the dsRNA-Seq reads. Thus, conventional sequencing was more sensitive than dsRNA-Seq at detecting the virus.

**Figure 3.**
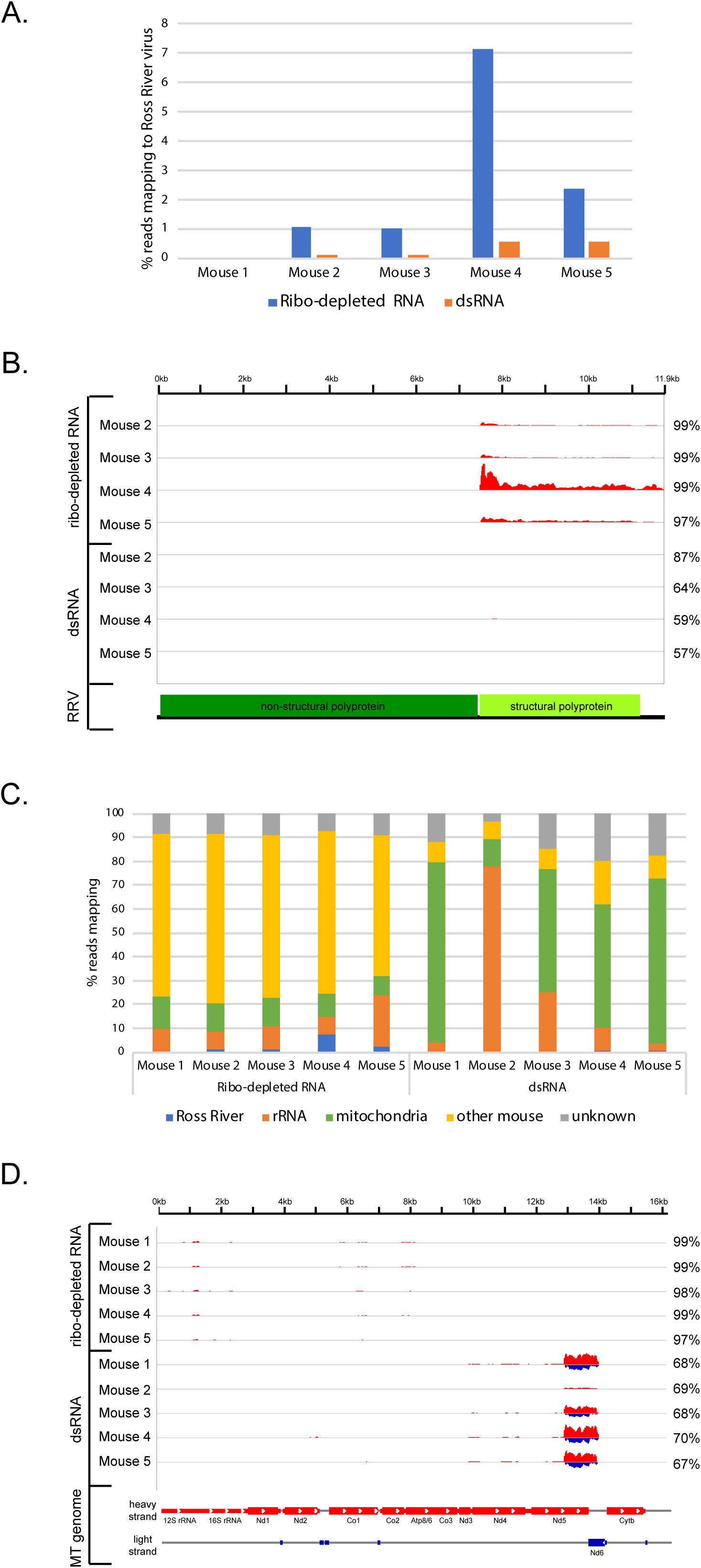
Comparison of dRNA-Seq and ribo-depleted RNA-Seq in identifying virus infections. (a) Percent of reads from ribo-depleted RNA-Seq libraries and dsRNA-Seq libraries prepared from mouse tissue samples that mapped to Ross River virus genome; (b) Bedgraphs indicating the base count of reads from RNA-Seq libraries and dsRNA-Seq libraries that aligned to each position of the Ross River virus genome. The bedgraphs were scaled relative to the total number of reads in each library and then the entire set was scaled identically as a group. Red indicates counts on positive strand. Blue indicates counts on negative strand. The percentage of the total number of reads that mapped to the Ross River virus that aligned to the positive strand are indicated on the right. A map of the Ross River virus with the location of the regions encoding the non-structural and structural polypeptides is indicated below; (c) Percentage of reads from ribo-depleted RNA-Seq libraries and dsRNA-Seq libraries from each mouse sample that mapped to the Ross River virus genome, mouse ribosomal sequences, mouse mitochondria genome, other mouse genome sequences or did not map (unknown); (d) Bedgraphs indicating the base count of reads from ribo-depleted RNA-Seq libraries and dsRNA-Seq libraries that aligned to each position of the mouse mitochondrial genome. The bedgraphs were scaled relative to the total number of reads in each library and then the entire set was scaled identically as a group. Red indicates counts on positive strand. Blue indicates counts on negative strand. The percentage of total reads in each sample that mapped to the mouse mitochondrial genome that aligned to the positive strand are indicated on the right. A map of the mouse mitochondrial (MT) genome is indicated below with the genes located on the heavy strand indicated in red and the genes on the light strand indicated in blue.

One reason that the Ross River virus sequences were depleted in the dsRNA is because the negative strand of the virus was in low abundance compared to the positive strand in the starting RNA population. Greater than 97% of the ribo-depleted reads that mapped to Ross River virus were derived from the positive strand of the virus (Figure 3B), and the mapping pattern was consistent with most reads deriving from the highly expressed subgenomic RNA which encodes the viral structural proteins (Figure 3B) [33-34]. Reads that cover the negative strand of the virus were present in the ribo-depleted datasets, though at very low abundance relative to the positive strand and within the error rate of the strand-specificity of the library (see Methods). Thus, while the overall abundance of Ross River virus sequences was reduced in the dsRNA-Seq datasets due to the removal of the abundant single-strand positive strand viral RNA during the dsRNA purification, the negative strand of the virus is well represented in the dsRNA-Seq reads (Figure 2C), allowing for the conclusion that the virus replicated in the tissues from which the RNA samples were derived.

A second explanation for the depletion of viral RNA sequences in dsRNA reads relative to the ribo-depleted RNA is that dsRNA-Seq enriches for host mitochondrial dsRNA. In all the dsRNA-Seq datasets but the Mouse 2 sample, 50% or more of the dsRNA reads map to the mitochondrial genome (Figure 3C). In contrast, mitochondrial sequences make up only 7-14% of the ribo-depleted RNA datasets (Figure 3C). The vast majority of the mitochondrial reads in the dsRNA-Seq samples mapped to both strands of a ∼1.2kb region of the mitochondrial (MT) genome, which corresponds to the position of the Nd6 gene (Figure 3D). The Nd6 gene is transcribed from the opposite strand of the mitochondrial genome compared to the other mitochondrial protein-encoding and rRNA genes (Figure 3D). Therefore, the enrichment of mitochondrial sequences in the dsRNA datasets is most likely due to the presence of sense/antisense mitochondrial transcripts from this region.

In summary, although dsRNA-Seq did not lead to enrichment of Ross River viral sequences, it provided evidence for viral replication which is a potential advantage of dsRNA-Seq. In the case of viruses producing subgenomic RNAs, such as alphaviruses, the presence of these RNAs in conventional RNA-Seq also provides evidence for viral activity. However, dsRNA-Seq is advantageous for the many viruses that only produce genomic length RNA. One way of improving dsRNA-Seq would be to develop ways to deplete the mitochondrial dsRNA sequences, effectively enriching for viral sequences.

### 3.4. dsRNA-Seq detects RNA viruses of multiple genome types in infected animals

Since we successfully detected RNA viruses in laboratory infected animals, we asked if dsRNA-Seq could do so in naturally infected animals. We obtained nine samples of total RNA isolated from various tissues of infected green tree python (*Morelia viridis*) (lung and a mixed lung/esophagus sample), rough scaled python (*Morelia carinata*) (lung), boa constrictor (*Boa constrictor*) (kidney), veiled chameleon (*Chamaeleo calyptratus*) (mixed lung/trachea/oral mucosa and two samples of mixed lung/liver/kidney), and mule deer (*Odocoileus hemionus*) (samples from brain and lymph node). These animals were found to be infected via various methods. The green tree and rough scaled pythons died of respiratory disease and were PCR positive for Morelia viridis nidovirus. Standard RNA-Seq confirmed the presence of a snake reptarenavirus and paramyxovirus in the boa constrictor. RNA-Seq was also used to detect a nidovirus coinfection in the chameleon samples (chameleon also died of respiratory disease). The mule deer was diagnosed with meningoencephalitis on postmortem exam, and was found to be infected with Caprine herpesvirus via DNA-Seq.

We used the same method and platform for dsRNA-Seq to sequence dsRNA isolated from 10 μg of total RNA from each sample. To screen for contaminants, we also processed a negative control sample containing water rather than RNA through the entire dsRNA purification and library preparation procedure. For each sample we obtained ∼10-29 million reads (Table S3).

To mimic a situation where host genomes were unavailable, we skipped host read filtering and directly assembled all reads into dsRNA contigs of 750 bases or longer. We then used the strand specific information to determine which contigs were assembled from dsRNA. Three to 87% of contigs were single-stranded possible contaminants, leaving 35 to 1013 contigs derived from dsRNA to analyze per sample, ranging from 0.5 to 23 kb (Table S3). To determine if the dsRNA contigs were derived from known or related viruses, we used BLASTn and BLASTx to search for similarity between the contigs at the nucleotide and protein level within the entire NCBI nt and nr databases.

Our analysis revealed two general points about the contigs from these samples (Figure 4 and Table S3). First, a number of dsRNA contigs showed similarity to bacterial sequences, many of which were also found in the negative control and are likely contaminants. Second, in general, the approach depleted most host sequences, with the exception of some mitochondrial dsRNA contigs in the deer samples, which is consistent with overlapping transcription in mammalian mitochondria producing some dsRNA.

**Figure 4.**
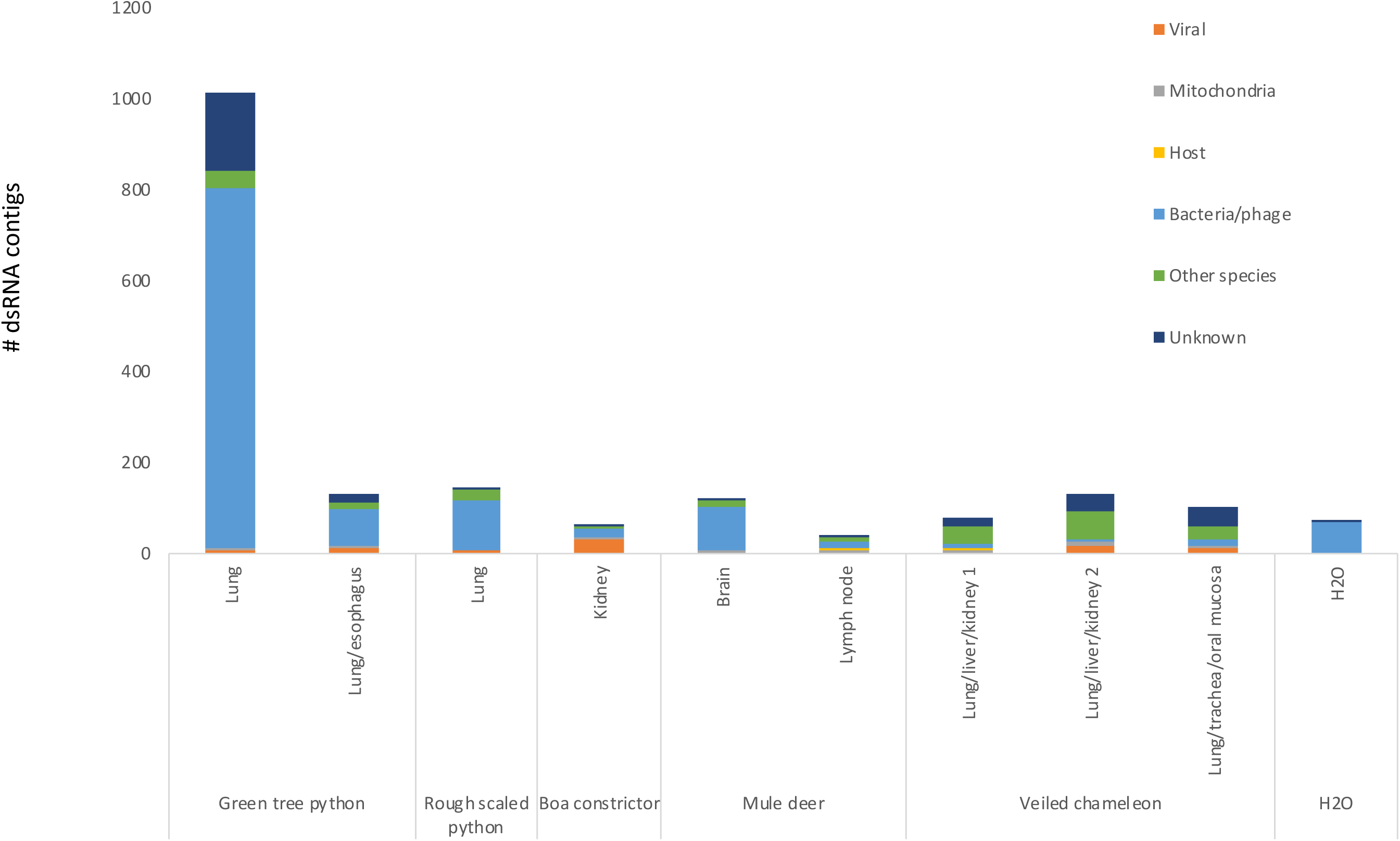
Classification of dsRNA contigs assembled from dsRNA isolated from reptiles and mule deer. Classification of dsRNA contigs from snake, mule deer, and chameleon samples according to BLASTn and BLASTx analysis. Viral, contigs that potentially infect the samples; host and mitochondria, contigs with hits to host nuclear or mitochondrial genomes, respectively; bacteria/phage, contigs with hits to bacterial or bacteriophage sequences; other species, contigs with hits to non-host eukaryotic organisms or viruses known to infect non-reptilian species; unknown, contigs without hits to known sequences.

More importantly, we were able to identify RNA viruses in the samples from the snakes and chameleon (Table 1). First, we identified Morelia viridis nidovirus (strain S12-1323) in the green tree and rough scaled python samples. This nidovirus is a single stranded, positive sense ∼32.4 kb RNA virus known to cause respiratory disease in pythons [21, 37]. Five contigs in the rough scaled python sample and eighteen contigs from the green tree python samples shared >90% sequence similarity on the nucleotide level to Morelia viridis nidovirus, and we were able to assemble partial genomes (Figure 5A). Since both strands of the viruses were present in the dsRNA-Seq reads, we can conclude these viruses were actively replicating in the animals.

**Figure 5.**
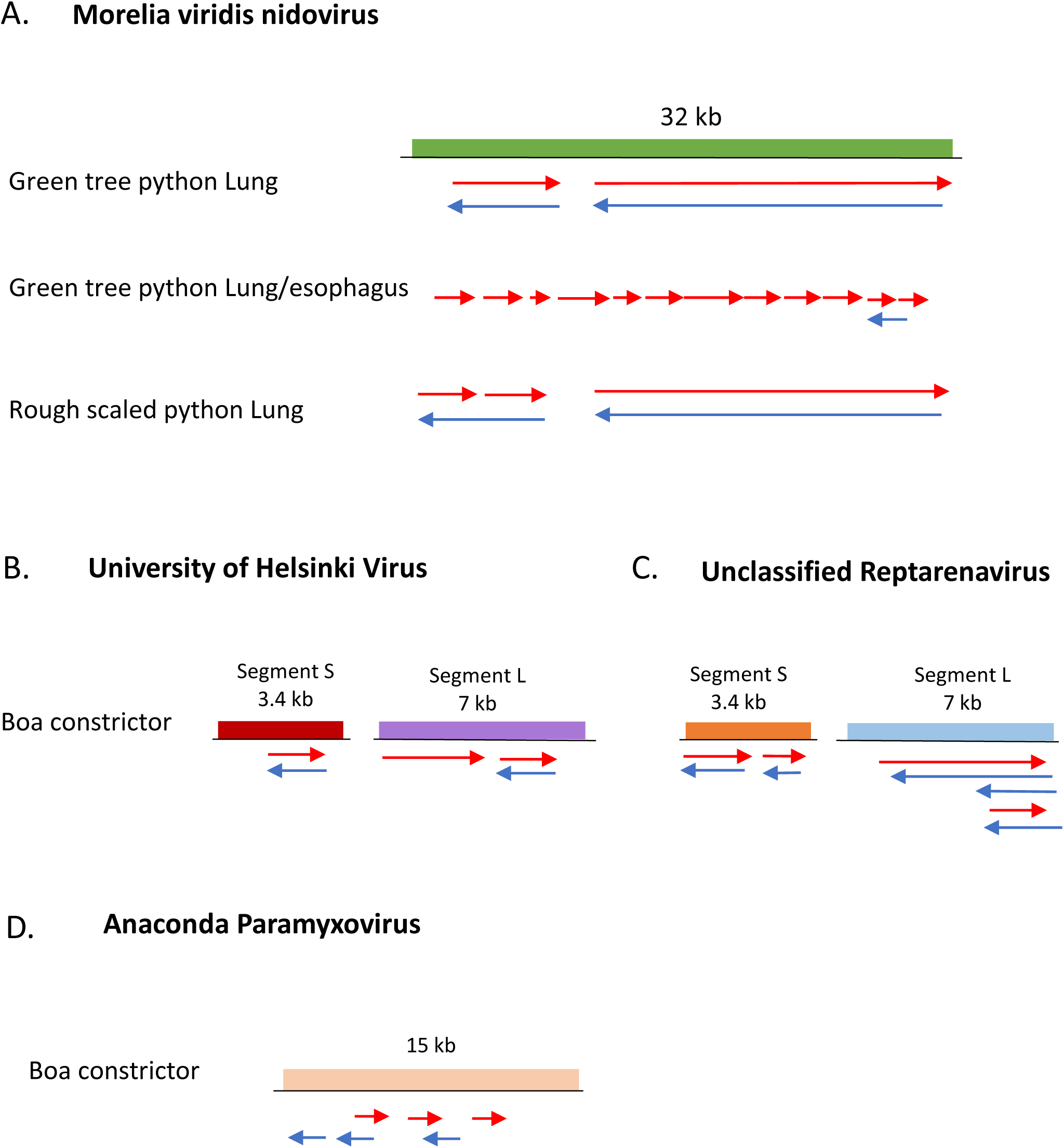
Snake dsRNA contigs aligned to viral genomes. Arrows indicate relative location of nucleotide sequences in contigs assembled from dsRNA that align with the corresponding virus. Contigs representing the positive strand of the virus are in red. Contigs representing the negative strand in blue. Longest region of contiguous similarity is shown. (a) dsRNA contigs isolated from green tree and rough scaled python lung and lung/esophagus pooled tissue mapped to Morelia viridis nidovirus genome (32 kb), represented by green boxes; (b) Boa constrictor dsRNA contigs mapped to University of Helsinki reptarenavirus segments (segment S ∼3.4 bp, segment L ∼7 kb), represented by orange boxes; (c) Boa constrictor dsRNA contigs mapped to Unidentified Reptarenavirus strain Reptarenavirus/Boa constrictor/California/snake38/2009 (segment S ∼3.4 kb, segment L ∼ 6.9 kb); (d) Boa constrictor dsRNA contigs mapped to Anaconda paramyxovirus isolate 1110RN047 (∼15 kb).

**Table 1.**
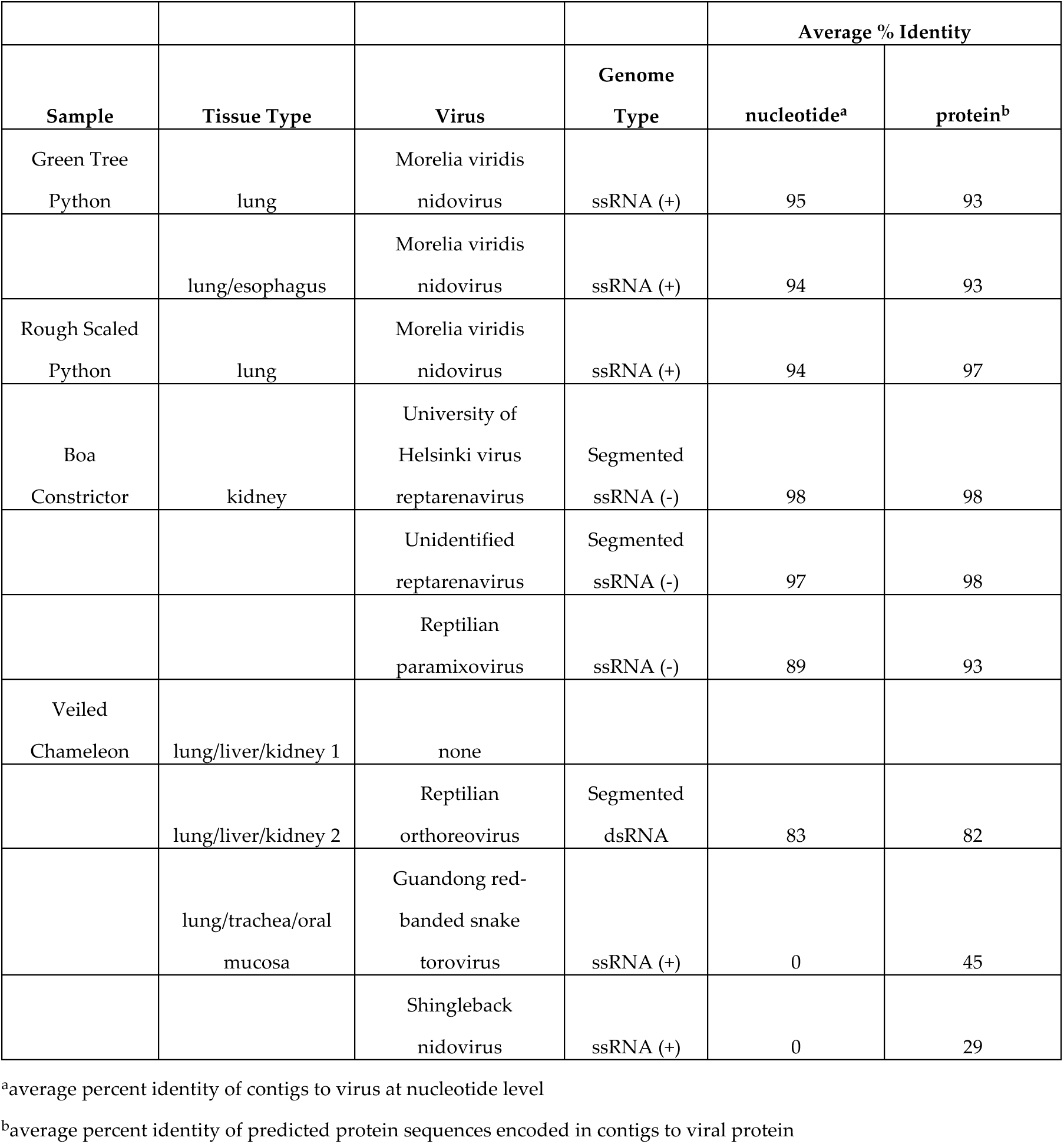
Viruses identified in reptile samples.

In the boa constrictor tissue we uncovered a coinfection of two distinct reptarenaviruses as well as a reptilian paramyxovirus. Reptarenaviruses contain negative-sense RNA genomes divided into two segments, a small (S ∼3.5 kb) and large (L ∼7 kb), and co-infections of this type of virus are reportedly common [38-39]. The S segment encodes the glycoprotein precursor (GPC) and the nucleoprotein (NP), whereas the RNA-dependent RNA polymerase (RdRp) and the Z protein (ZP) are encoded by the L segment [38]. Two contigs in this sample were >98% identical at the nucleotide level to University of Helsinki Virus and mapped to portions of both segments of this reptarenavirus (Figure 5B). Fourteen boa constrictor contigs were >90% identical on the nucleotide level and seven additional contigs were >96% identical on the protein level to previously sequenced but unclassified reptarenaviruses (Figure 5C and Table S3). These contigs might represent multiple viruses. We identified contigs that mapped to full S and L unclassified reptarenavirus segments.

We also found four contigs in the boa constrictor sample that were >90% identical at the nucleotide level and three additional contigs at the protein level to a number of reptilian paramyxoviruses. (Figure 5D and Table S3). Paramyxoviruses belong to the family *Paramyxoviridae* and are negative sense, single stranded viruses associated with neuro-respiratory disease in reptiles [40]. We were not able to assemble a complete paramyxovirus genome, although we obtained hits for genes encoding the nucleoprotein, fusion protein, hemagglutinin-neurimidase, and RNA-dependent RNA polymerase (L) (Figure 5D). Both the reptarenaviruses and the paramyxovirus had dsRNA-Seq reads that corresponded to both strands of the virus, indicating they had replicated in this boa constrictor.

In addition to confirming viruses in snakes that were previously found via standard RNA-Seq, dsRNA-Seq allowed us to detect a dsRNA virus in chameleon tissue that was not detected by RNA-Seq. Three contigs in the pooled lung/liver/kidney sample 2 were 76-87% identical on the nucleotide level to a reptilian orthoreovirus, within the family *Reoviridae*, a segmented, dsRNA linear virus that contains ten segments coding for 12-13 proteins. Twelve additional contigs were 47-95% identical on the protein level to proteins encoded by the same virus (Figure 6A). dsRNA-Seq enriched for this dsRNA virus given that no reads which align to the orthoreovirus contigs were detected in RNA-Seq data obtained from the same animal. In addition, we uncovered a possible coinfection of two nidoviruses. Nine contigs in the chameleon pooled lung, trachea, and oral mucosa sample were 28-59% identical at the protein level to a 7.5 kb Guangdong red-banded torovirus protein coding sequence (Figure 6B). Two additional contigs shared 26-29% identity to a small region of the protein coding sequences of Shingleback nidovirus (Figure 6C). Given their low similarity to known nidoviruses, these contigs likely represent previously uncharacterized nidoviruse(s).

**Figure 6.**
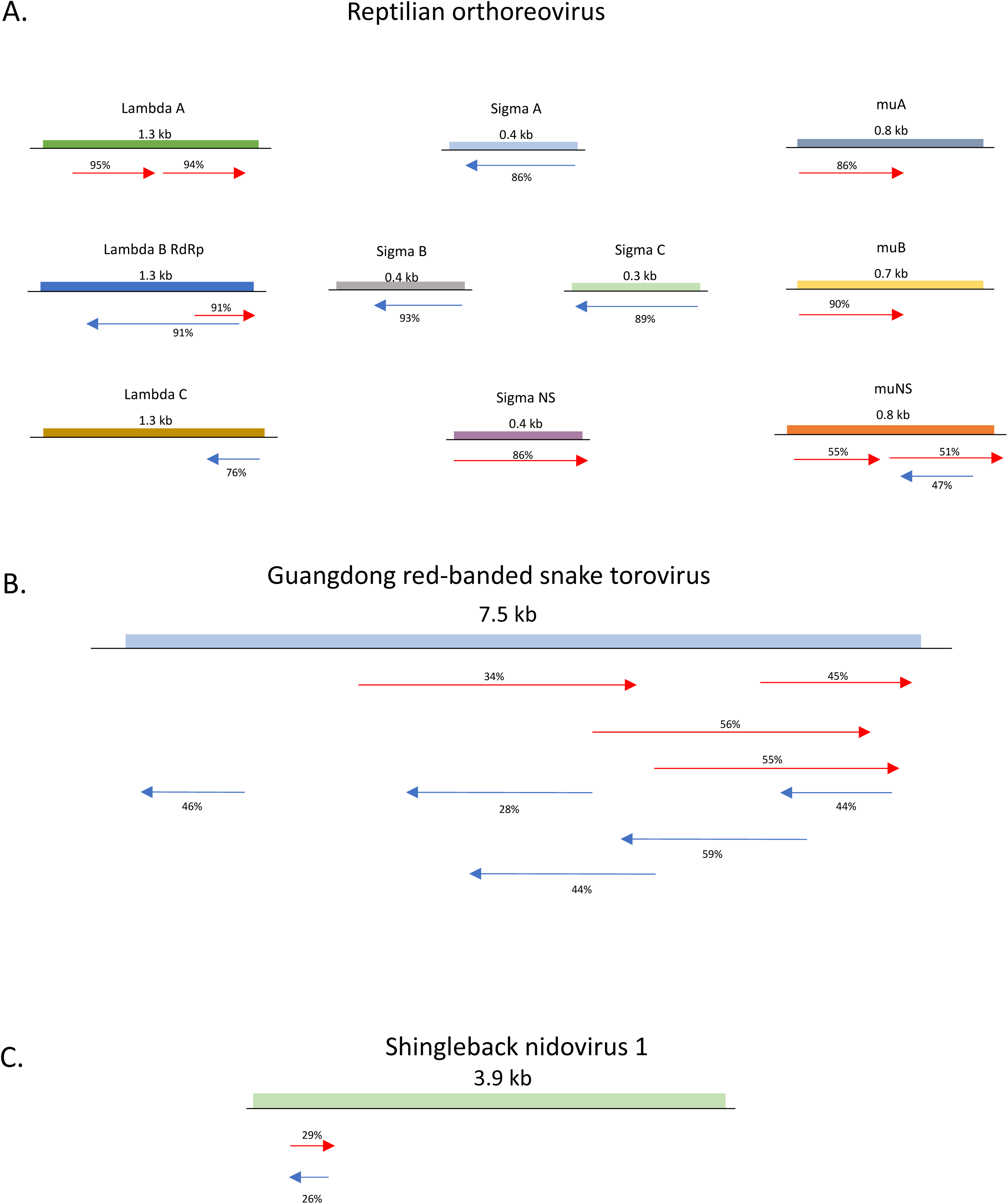
Chameleon dsRNA contigs with protein similarity to known viruses. The length of viral genomes or genome segments are indicated. Coding regions are indicated by colored boxes. The locations where contigs shared protein similarity to the virus are indicated by arrows, Contigs representing the positive strand of the virus are displayed by red arrows; the negative strand by blue arrows. The percent identity between the region in the contig and the corresponding protein coding sequence in the reference virus is indicated above the arrow. (a) Schematic of chameleon dsRNA contigs aligned to portions of all ten protein coding segments of reptilian orthoreovirus; (b) Schematic displaying chameleon dsRNA contigs aligned to Guangdong red-banded snake torovirus; (c) Schematic displaying chameleon dsRNA contigs aligned to Shingleback nidovirus.

We did not identify any viral contigs in the two mule deer samples (brain and lymph node tissue.) Previous DNA-Seq [41] detected a herpesvirus in the mule deer. Herpesvirus is a dsDNA virus, which likely explains our inability to detect viral reads via dsRNA-Seq.

We used the boa constrictor sample where we had both dsRNA-Seq and standard RNA-Seq data to examine the sensitivity of the two methods. We mapped the boa constrictor reads from both methods to the assembled unclassified reptarenavirus contigs and found that 10.16% of standard RNA-Seq reads aligned, compared to 42.25% of dsRNA-Seq reads. We performed the same analysis with the University of Helsinki virus contigs and found that 1.32% of standard RNA-Seq reads aligned compared to 5.28% of dsRNA-Seq reads. For the Anaconda paramyxovirus contigs, 0.05% of standard RNA-Seq aligned, compared to 0.02% of dsRNA-Seq. These results indicate that dsRNA-Seq can be more sensitive than standard RNA-Seq in detecting viruses in naturally infected animals, however this varies for different viruses.

To elucidate the efficacy of rRNA depletion by dsRNA-Seq, we compared the percentage of boa constrictor standard RNA-Seq reads and dsRNA-Seq reads that aligned to boa constrictor imperative mitochondrial rRNA sequences (AM236348.1) which are available at NCBI. 16.84% of standard RNA-Seq boa constrictor reads aligned to rRNA, while only 0.47% of dsRNA-Seq reads aligned to rRNA. The 36-fold reduction in rRNA demonstrates the usefulness of dsRNA-Seq for sequencing organisms without commercially available rRNA depletion reagents.

In summary, dsRNA-Seq identified viruses of various RNA genome types in naturally infected animals and allowed us to assemble partial and full-length genomes of viruses infecting snake and chameleon tissue.

## 4. Discussion

By several criteria the two-step method we have developed for purifying dsRNA from total RNA samples is very effective. For example, in the mouse samples, we observe strong depletion of ribosomal RNA and high enrichment of host dsRNA in the dsRNA-Seq libraries. Total RNA from mammalian cells is ∼85% rRNA and ∼15% mRNA and other RNA species. Based on western blot analysis of total RNA using the anti-dsRNA antibody, we estimated that only ∼0.01 to 0.1% of total RNA isolated from mammalian tissue was dsRNA. Given these ratios, if the dsRNA purification method was 99.9% effective at removing rRNA and host single-stranded RNA we would expect ∼42% of the reads in the purified dsRNA to be rRNA and ∼50% being host dsRNA. In all the dsRNA-Seq samples, except for the sample from mouse 2, we observed an even stronger depletion of ribosomal sequences (Fig 3C) indicating that removal of rRNA was very effective (the nuclease treatment and/or anti-dsRNA immune-purification appears to have been less effective in the mouse 2 sample than the remaining samples). Moreover, there is considerable enrichment of host dsRNA in the dsRNA-Seq libraries with at least 50% of the dsRNA reads arising from transcripts from opposite strands of the mitochondrial genome (Fig 3C and D). In addition, a global analysis of the alignment of the ribo-depleted and dsRNA-Seq reads to known mouse transcripts revealed that of the dsRNA-Seq reads that aligned to known mouse transcripts, ∼50% were derived from the forward strand and ∼50% from the reverse strand (data not shown). In contrast, 90% of the ribo-depleted RNA-Seq reads prepared using the same strand-specific library kit mapped to the forward strand of the known transcripts (data not shown). This observation is consistent with both strands of the mouse transcripts being present in the purified dsRNA before the strand-specific library was prepared. Isolation of dsRNA by several different strategies has been used to detect viruses in plants [42-43], fungi [44] and microbial communities [45]. Our method is likely to be more selective because it employs a two-step purification scheme.

We posited that dsRNA-Seq should be able to detect all types of RNA viruses given that dsRNA has been detected in cells infected with positive, negative and ambisense single-stranded RNA viruses as well as dsRNA viruses [46-47]. Using dsRNA-Seq we successfully identified and assembled full or partial genomes for non-segmented and segmented negative- and positive-sense RNA viruses, dsRNA viruses, and as well as ambisense RNA viruses. In all cases, dsRNA-Seq provided evidence that the viruses had replicated in the tissue examined by detecting the positive and negative strand sequences of each virus.

This method is useful to distinguish actively replicating RNA viruses from non-replicating or contaminating viral sequences by revealing the presence of both viral strands. It should be noted however that this method does not necessarily eliminate viruses as potential pathogens since some viruses may replicate at too low a level to be detected or may not be actively replicating at the time of sampling. One disadvantage of dsRNA-Seq is that it cannot be used to detect single-stranded RNA viruses from clinical samples where viruses would primarily only be present as viral particles, such as serum or cerebral spinal fluid. dsRNA-Seq is particularly useful for sequencing viruses from organisms without sequenced genomes and/or with no commercially available rRNA depletion reagents. Another advantage of this approach over standard sequencing methods is that it may increase the ability to detect dsRNA viruses, such as the dsRNA orthoreovirus we identified in chameleon tissue.

Although we identified viral contigs assembled from purified dsRNA by searching for similarity between the contigs and known viral sequences, it would have been possible to recognize these contigs as being potential viruses or viral segments without the contigs sharing similarity either on the nucleotide or protein level with a known virus. First, these contigs were likely not derived from the host since they did not map to the host nuclear or mitochondrial genome. Second, each of the contigs encode open reading frames that span across nearly the entire length of the contig similar to the genomic organization of most eukaryotic RNA viruses or viral segments. Assembly of long viral contigs is critical for the recognition of viruses that are highly divergent from previously described viruses, such as the chameleon orthoreovirus and nidoviruses, because it increases the ability to detect remote similarities. Thus, it is theoretically possible using dsRNA-Seq to identify potential novel or synthetic viruses that have limited to no sequence similarity to known viruses.

It should be noted that sequences present as “dsRNA” in dsRNA-Seq libraries are not necessarily base-paired with each other in vivo. Hybrids between sense and antisense species during or post RNA isolation would also be expected to be purified as dsRNA. During RNA viral replication and transcription, strands may be separated to limit detection of dsRNA by the host innate immune system. However, the presence of both viral strands in total RNA should be sufficient to allow recovery of viral sequences in the dsRNA. Consistent with this idea, there were more dsRNA reads that mapped to the region of Ross River virus which corresponded to the abundant subgenomic RNA (Fig 2C) than to other regions of the virus, presumably because the high abundance of the subgenomic RNA increased the likelihood of there being hybrid molecules of positive and negative strands in this region.

## Author Contributions

Conceptualization, R.P. and C.J.D.; Data Curation, C.J.D and H.R.S.; Formal Analysis, C.J.D, H.R.S. and L.L.H-H.; Funding Acquisition, R.P., E.M.P. and S.L.S.; Investigation, C.J.D, H.R.S. and L.L.H-H; Methodology, C.J.D and H.R.S. Project administration, R.P. and C.J.D.; Resources, A.C.S., S.L.S., K.C.H., T.E.M., L.L.H-H, M.D.S., J.H.M. and E.M.P; Supervision, R.P; Visualization, C.J.D and H.R.S.; Writing-Original Draft Preparation, C.J.D, H.R.S. and R.P.; Writing-Review and Editing, C.J.D, H.R.S., L.L.H-H, M.D.S., J.H.M., E.M.P., T.E.M., S.L.S. and R.P.

## Acknowledgments

We thank the Next-Gen Sequencing Core Facility at the University of Colorado, BioFrontiers Institute, which performed the Illumina sequencing and shared helpful information regarding low input library kit selection. This research was conducted using the resources of the BioFrontiers Computing Core at the University of Colorado, BioFrontiers Institute. We thank the BioFrontiers Computing Core staff for their help and support.

## Supplementary Materials

**Figure S1.**
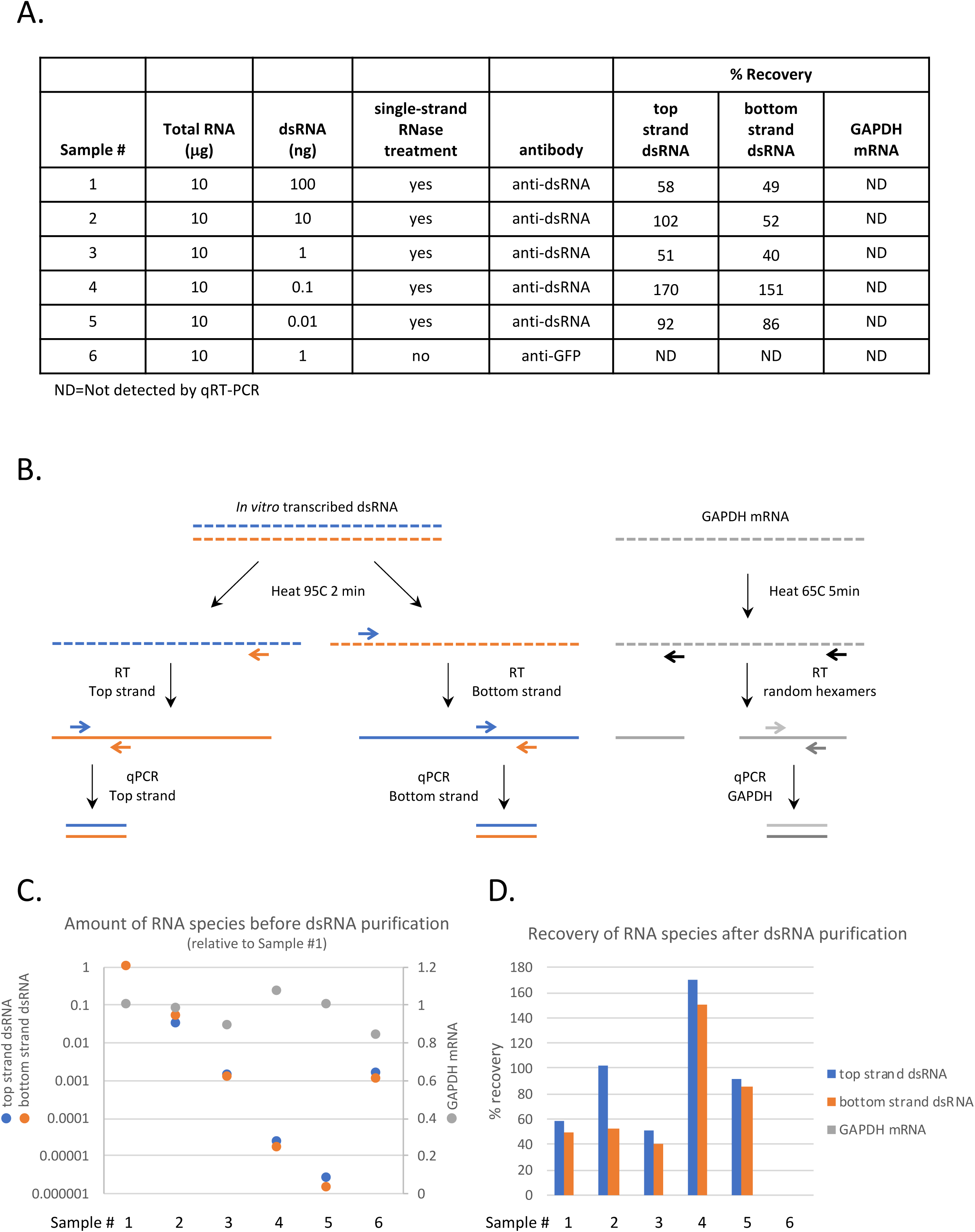
Analysis of dsRNA purification method. (a) The two-step dsRNA purification method described in Materials and Methods was used to purify dsRNA from samples containing a fixed amount of total RNA isolated from human tissue culture cells spiked with varying amounts of a 0.9kb *in vitro* transcribed dsRNA [45]. The RNA samples were treated with or without single-strand specific RNase and then dsRNA was isolated using either 5mg of the anti-dsRNA antibody J2 or a control antibody against GFP. (b) Schematic of the three qRT-PCR reactions used to determine the amount of the individual strands of the dsRNA or a single-strand RNA (GAPDH mRNA) in the samples before and after dsRNA purification. RNA indicated by dashed lines, DNA by solid lines. (c) Amount of the three RNA species in starting samples before dsRNA purification procedure plotted relative to Sample 1. Relative amount of dsRNA strands reported on left axis on log scale. Relative amount of GAPDH mRNA reported on right axis on linear scale. (d) Recovery of the three RNA species after dsRNA purification. In all samples, the recovery of GAPDH mRNA was below the detection limits of the qRT-PCR.

**Figure S2.**
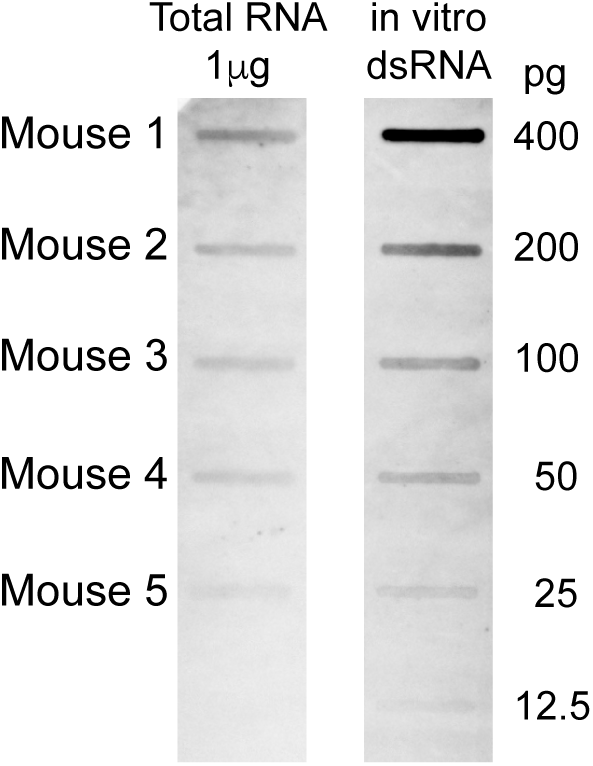
Anti-dsRNA western analysis of mouse total RNA. Antibody against dsRNA was used to estimate the amount of dsRNA in total RNA isolated from mouse tissue samples. 1μg of total RNA from each mouse total RNA sample was loaded in a slot blot and compared to a dilution series of a known amount of in vitro transcribed 0.9kb dsRNA [45]. Anti-dsRNA immunoblotting was performed as described in [17, 45].

**Figure S3.**
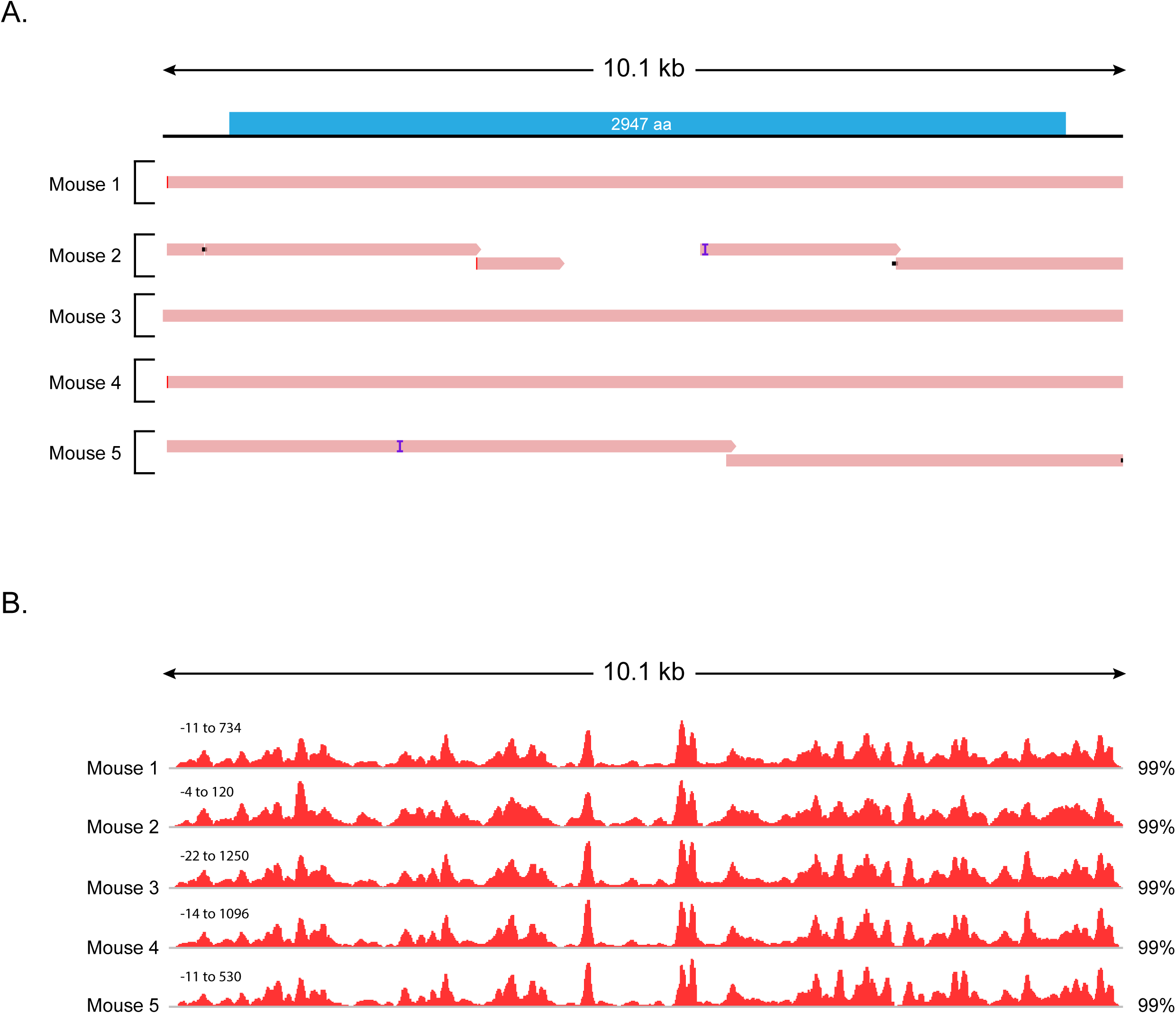
Picorna-like viral contigs assembled from mouse dsRNA-Seq libraries are derived from single-stranded RNA. (a) Alignment of contigs representing novel picorna-like virus. The longest contig with similarity to Sanxia picorna-like virus 7 (MM3_TRINITY_DN634_c0_g1_i5) was selected as a representative. Contigs assembled from each mouse dsRNA-Seq library were aligned to the representative contig using bwa-mem. The location of the long open reading frame present in the representative contig is illustrated at the top. (b) Bedgraphs indicating the base count of reads from dsRNA-Seq libraries that aligned to each position of the representative picorna-like virus contig. Red indicates counts on positive strand. Blue indicates counts on negative strand. The percentage of total reads that mapped to the representative picorna-like virus contig that aligned to the positive strand are indicated on the right. The bedgraphs were not scaled relative to total number of dsRNA-Seq reads in each sample. Each read set was scaled individually, scale is indicated at top left for each set.

**Table S1. Summary of contigs assembled from dsRNA-Seq reads from Vero cell culture samples.** Includes summary of dsRNA read mapping, summary of the classification of the contigs and lists of the contigs from each sample including their length, total number reads mapped, percentage of reads mapped to forward strand, and the classification of the contig.

**Table S2. Summary of double stranded contigs assembled from dsRNA-Seq reads from mouse tissue samples.** Includes summary of dsRNA read mapping, summary of the classification of the contigs and lists of the contigs from each sample including their length, total number reads mapped, percentage of reads mapped to forward strand, and the classification of the contig.

**Table S3. Summary of dsRNA contigs assembled from dsRNA-Seq reads from reptile and mule deer tissue samples.** Includes summary for dsRNA sequencing, summary of the classification of the contigs and lists of the contigs from each sample including their length, total number reads mapped, percentage of reads mapped to forward strand, and the classification of the contig.

File S1. Vero 1 contigs fasta file

File S2. Vero 2 contigs fasta file

File S3. Vero 3 contigs fasta file

File S4. Mouse 1 contigs fasta file

File S5. Mouse 2 contigs fasta file

File S6. Mouse 3 contigs fasta file

File S7. Mouse 4 contigs fasta file

File S8. Mouse 5 contigs fasta file

File S9. Green Tree Python Lung dsRNA contigs fasta file

File S10. Green Tree Python Lung Esophagus dsRNA contigs fasta file

File S11. Rough Scaled Python Lung dsRNA contigs fasta file

File S12. Boa Constrictor Kidney dsRNA contigs fasta file

File S13. Veiled Chameleon Lung Trachea Oral Mucosa dsRNA contigs fasta file

File S14. Veiled Chameleon Lung Liver Kidney 1 dsRNA contigs fasta file

File S15. Veiled Chameleon Lung Liver Kidney 2 dsRNA contigs fasta file

File S16. Mule Deer Brain dsRNA contigs fasta file

File S17. Mule Deer Lymph Node dsRNA contigs fasta file

File S18. Negative Control dsRNA contigs fasta file

